# Substrate chemistry in cancer bioelectronics: phenotype-dependent cell-electrode coupling and motility

**DOI:** 10.64898/2026.10.21.755265

**Authors:** Catarina F. Jones, Brindha Jeyaseelan, Rosalia Moreddu

## Abstract

Electrode coatings are routinely applied to promote cell attachment in bioelectronics, yet their influence on what the electrode reports is less examined. This is particularly relevant for cell types whose electrical signatures are yet to be fully understood and where coating selection is largely inherited from neuroscience. Here, we compare three breast epithelial phenotypes of increasing invasiveness on microelectrodes carrying six surface chemistries: uncoated gold, poly-L-lysine (PLL), fibronectin, laminin, and PLL under-layered with each protein (PLL+F, PLL+L). Continuous impedance recording at 1 kHz and 41.5 kHz over the first 17 h, spatial impedance mapping at 24 h, 50 h migration assays and single-cell trajectory analysis were used to evaluate cell–electrode coupling and relate it to motility in the two extremes of the spectrum. Each cell line produced its electrical signature, and the effect of coating depended on phenotype. Initial coupling and motility were found to be inversely related in the invasive line and unrelated in the non-tumorigenic one. We conclude that the electrode coating is an experimental variable that interacts with phenotype and carries its own electrical contribution, and that the choice of coating must be driven, beyond physiological relevance, by the requirement for adherent vs. motile cells.

## Introduction

Cancer is increasingly understood as a disease with an electrical dimension. Cancer cells carry a depolarized resting membrane potential relative to their normal counterparts, and this shift regulates proliferation: forcing depolarization can induce cancer-like growth in otherwise normal tissue, while restoring polarization suppresses it^1,2^. Ion channels that set and modulate membrane potential are overexpressed in many solid tumors, to the point that cancer has been framed as a family of channelopathies^3^. In breast cancer specifically, the voltage-gated sodium channel Nav1.5 is enriched in some aggressive cell types, where it promotes migration and invasion^4^. These findings have already reached the clinic: tumor-treating fields, which disrupt mitosis with alternating electric fields, are an approved therapy for glioblastoma^5^, and irreversible electroporation is used to ablate pancreatic and hepatic tumors that are inaccessible to thermal methods^6^. The reverse direction, using electronics to read cancer cell state, is less mature but holds a complementary promise: a label-free, continuous and quantitative readout of phenotype that could support diagnosis, drug screening and biophysics studies^7^.

Electrical impedance spectroscopy is the most established route to such a readout. Since the demonstration that cells settling on microelectrodes alter its impedance in proportion to their coverage, spreading and junctional tightness^8^, electric cell–substrate impedance sensing (ECIS) has been used to follow attachment kinetics^9^, barrier formation^10^, cytotoxicity^11^ and electrically wounded monolayer recovery^12^. The frequency dependence of the signal allows the contributions of paracellular current, which dominates at low frequency, and of membrane capacitance and electrode coverage, which dominate at high frequency, to be separated^10^. Applied to cancer, whole-cell impedance spectroscopy distinguishes breast cell lines of different malignant potential^13^, ECIS resolves cancer cell adhesion, proliferation and drug response in real time^14^, and organic electrochemical transistors extend the approach to mechanically compliant electrodes capable of resolving barrier integrity with high sensitivity^15,16^. All these systems measure the degree to which cells couple to the electrode. Coupling, in turn, is governed by adhesion, and the latter depends on surface functionalization.

Electrodes are routinely coated to encourage attachment, and the choice of coating is typically inherited from neuroscience or general cell-culture practice rather than tested for its influence on the readout. Poly-L-lysine (PLL) is the default in much of the electrophysiology and impedance literature because it works for almost any cell type^17^. Yet PLL promotes adhesion electrostatically, through the attraction between its positive charges and the negatively charged glycocalyx and does not engage the integrin receptors through which cells normally bind the extracellular matrix^18^. Integrin engagement, in fact, induces focal adhesions as signaling hubs that convert substrate chemistry and mechanics into cytoskeletal tension, spreading, proliferation and migration^19^. Fibronectin and laminin are recognized by different integrin heterodimers and elicit different adhesive and migratory programs^20^. The transition from a cohesive epithelium to individually motile mesenchymal-like cells reorganizes the adhesive machinery that impedance measures: E-cadherin-based cell–cell junctions are lost, integrin-based cell–substrate contacts become dynamic, and migration shifts from collective to single-cell modes^21^. Therefore, the coating is part of the biological environment and affects the measurement.

Cells couple to an electrode differently and respond to a given coating differently, but whether and how they do so is poorly investigated. Breast cancer provides a well-characterized test model for such study. The non-tumorigenic MCF10A line forms polarized and junction-rich monolayers^22^, MDA-MB-468 cells are basal-A, E-cadherin-positive and grow in cohesive clusters, MDA-MB-231 cells are basal-B, E-cadherin-negative, spindle-shaped and highly invasive^23^. These lines span a gradient of epithelial-to-mesenchymal character within a single tissue of origin and have been compared electrically before^13^, but not across substrates. In this work, MCF10A, MDA-MB-468 and MDA-MB-231 cells were cultured on interdigitated gold microelectrodes carrying six surface chemistries (uncoated gold, PLL, fibronectin, laminin, and PLL under-layered with each protein), and were followed through four readouts that probe successive stages of the cell–electrode interaction: live/dead staining, continuous impedance at 1 kHz and 41.5 kHz over the first 17 h to capture attachment and spreading, spatial resistance mapping at 24 h to capture the organization of the layer, and 50 h migration assays with single-cell trajectory analysis to capture the functional consequence for motility (**Figure 1**).

**Figure 1.**
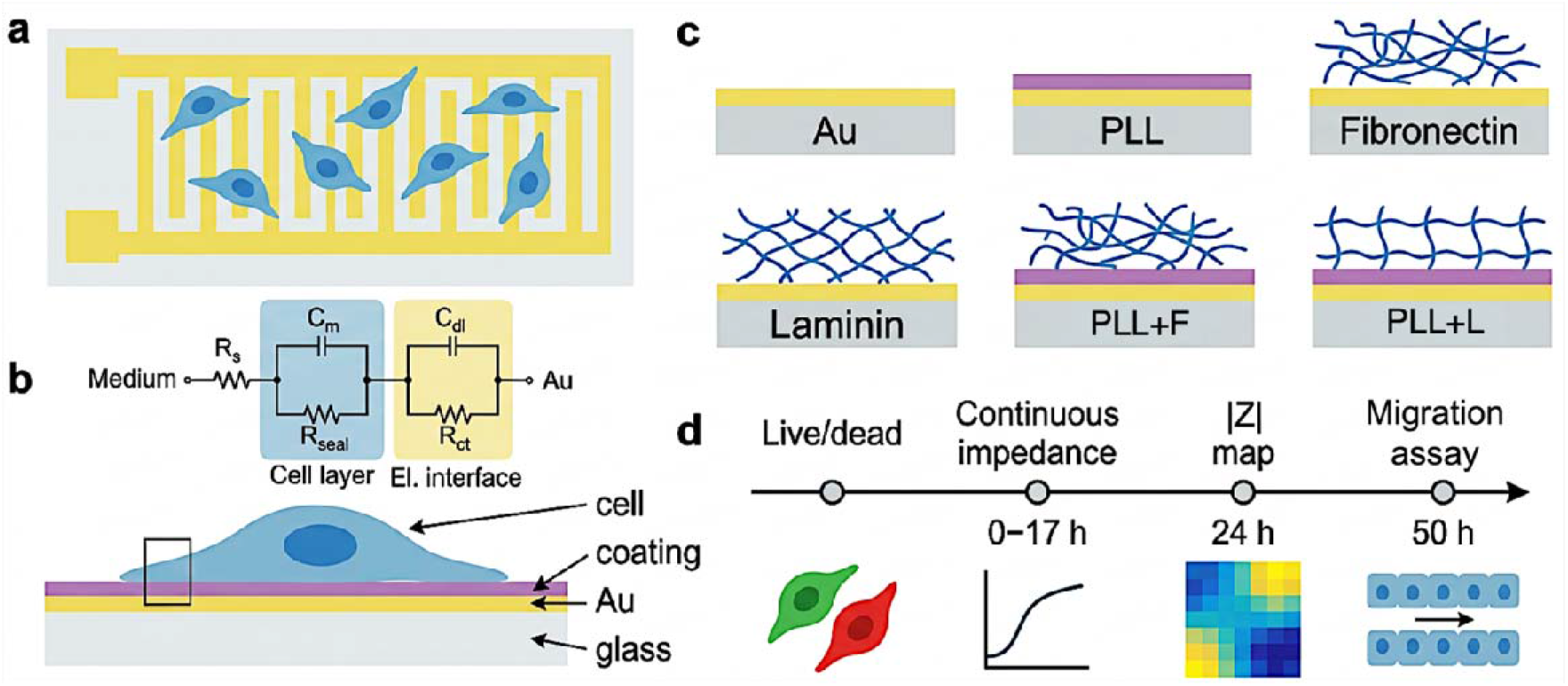
Experimental platform and workflow. **(a)** Plan view of the interdigitated gold microelectrode on a glass substrate with seeded cells. **(b)** Side view of the cell–electrode interface showing the cell, the coating (purple), the gold electrode and the glass substrate; the boxed region marks the interface described by the equivalent circuit above. Current from the medium passes through the solution resistance (R_s_), then across the cell layer either through the membrane capacitance (C_m_) or around and beneath the cells through the seal resistance (R_seal_), and finally through the electrode interface, modelled as the double-layer capacitance (C_dl_) in parallel with the charge-transfer resistance (R_ct_). **(c)** The six substrate chemistries used in this work. **(d)** Readouts applied to each cell line on each substrate: live/dead staining, continuous impedance recording at 1 kHz and 41.5 kHz over the first 17 h after seeding, spatial mapping of |Z| across the array at 24 h, and 50 h migration assays with single-cell trajectory tracking.

## Results

Three breast epithelial phenotypes were compared: the non-tumorigenic MCF10A line, the intermediate MDAMB468 line and the highly invasive MDAMB231 line. Each line was cultured on interdigitated gold microelectrodes or an array of 4×4 PEDOT: PSS microelectrodes carrying one of the six surface chemistries. Live/dead staining was performed on the coated electrodes to confirm cytocompatibility (**Figure 2**). On all surfaces and for all lines, the fields were vastly dominated by live (green) cells (**Figure 2a**). Quantification confirms viability close to 100% in every condition (**Figure 2b**). The MCF10A population carries a small fraction of dead cells on every surface, slightly larger on PLL+L, attributed to basal turnover within the monolayer^22^, which did not affect the subsequent coupling of this line, which was highest of the three on all substrates.

**Figure 2.**
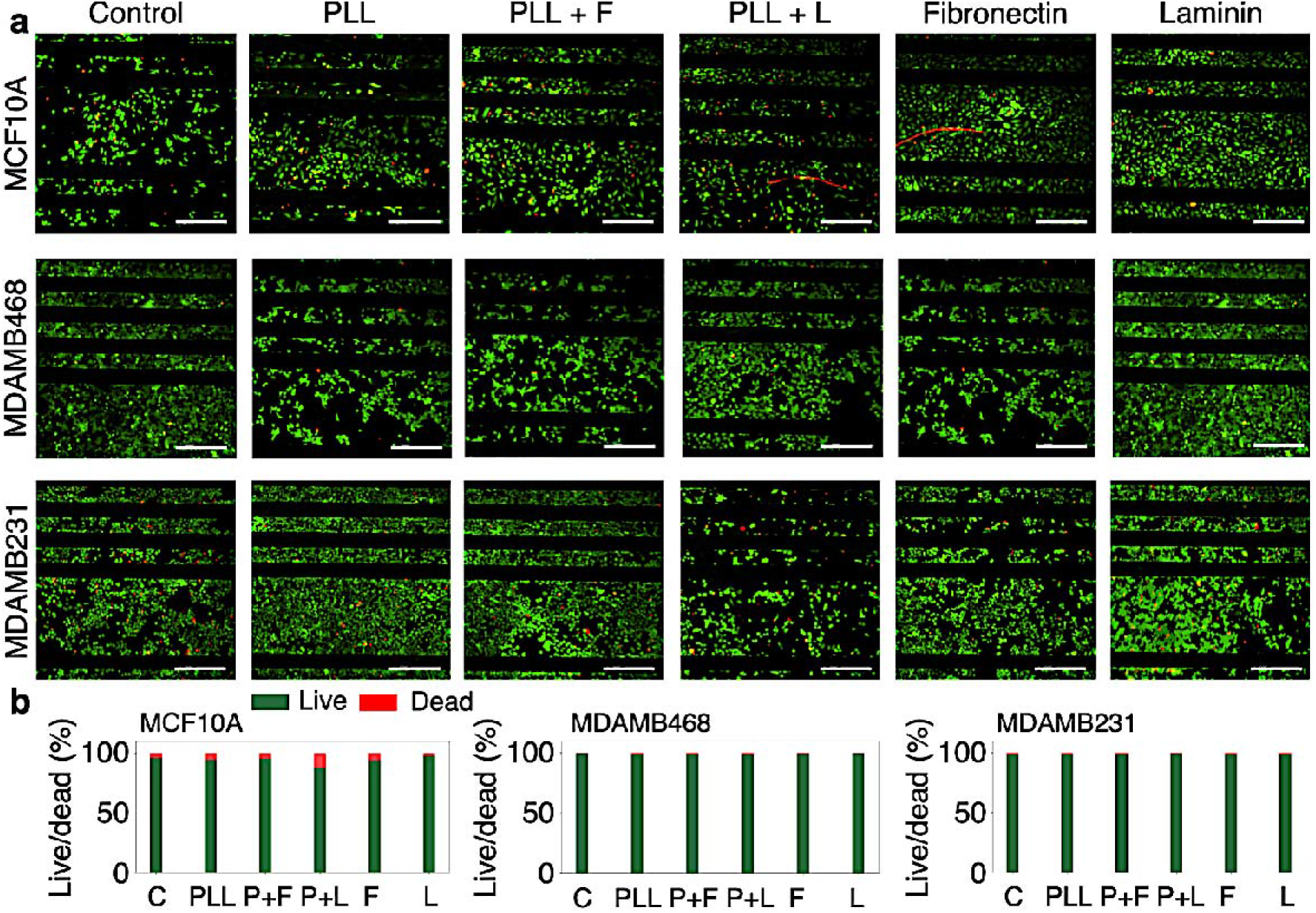
Substrate cytocompatibility across the tested breast cell phenotypes. **(a)** Live/dead fluorescence micrographs (live cells in green, dead cells in red) of MCF10A, MDAMB468, and MDAMB231 cells seeded on interdigitated microelectrodes across the six conditions. Scale bar: 200 µm. **(b)** Quantification of live and dead fractions (mean of n = 3 fields per condition).

### Time-resolved cell–electrode coupling

Steady-state impedance refers to the absolute |Z| at 1 kHz mapped across a separate array type at 24 h. The former reports attachment and spreading kinetics, the latter reports the organization of the settled layer superimposed on the coating background. Throughout this text, initial coupling refers to the changes in resistance and capacitance over the first 17 h after seeding, expressed relative to the coated electrode immediately before seeding, so that the offset given by the coating is removed at t = 0. Continuous electrical impedance spectroscopy was used to follow the first 17 h after seeding, the period over which cells sediment, attach and spread onto the electrode^24^ (**Figure 3**). Two frequencies were recorded. At 1 kHz the measurement is sensitive to current forced to pass around and between cells, and so reports on the tightness of cell–substrate and cell–cell contacts^25,26^. At 41.5 kHz the signal is dominated by the dielectric coverage of the electrode by the basal membrane, and so reports on spreading^26^. Both the capacitance change (ΔC) and the absolute impedance/resistance change (ΔR) were recorded at each frequency, since the two respond differently to the same event: coverage of the electrode by an insulating cell layer raises ΔR and lowers ΔC at the high frequency^9^. **Figure 3b–d** shows the channel most sensitive to each process (ΔR at 1 kHz, ΔC at 41.5 kHz). The complementary channels are in **Figure S1** and **Figure 3f–i**, and all four channels are compared for each substrate in **Figures S2–S4**. Endpoint values at 17 h are summarized in **Figure 3e** and **Table S1**.

**Figure 3.**
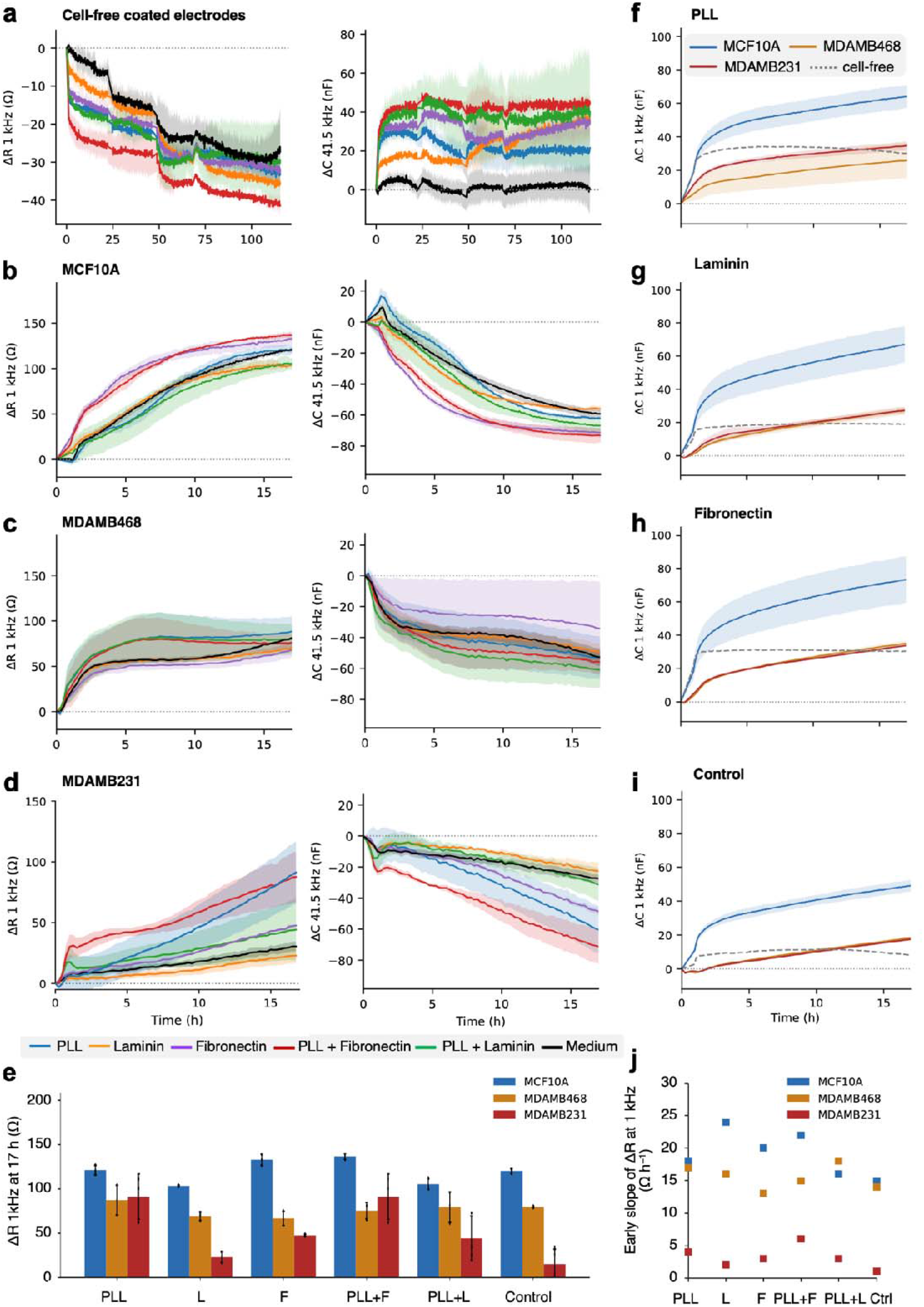
Time-resolved cell–electrode coupling across phenotypes and substrate coatings. **(a–d)** Continuous impedance spectroscopy on interdigitated gold microelectrodes carrying six surface chemistries: PLL (blue), laminin (orange), fibronectin (purple), PLL + fibronectin (red), PLL + laminin (green) and uncoated electrode (black). Left column: change in resistance at 1 kHz (ΔR, Ω), sensitive to current passing around and between cells. Right column: change in capacitance at 41.5 kHz (ΔC, nF), sensitive to dielectric coverage of the electrode by the basal membrane. The complementary channels (ΔR at 41.5 kHz, ΔC at 1 kHz) are shown in *Figure S1*. **(a)** Coated electrodes without cells monitored in DMEM for 120 h. Steps at 24, 48 and 70 h correspond to medium renewals. **(b–d)** Electrodes seeded with **(b)** MCF10A, **(c)** MDAMB468 and **(d)** MDAMB231 cells over the first 17 h after seeding. **(e)** ΔR at 1 kHz at 17 h for each line and substrate (mean ± SD). **(f–i)** ΔC at 1 kHz over 17 h for MCF10A (blue), MDAMB468 (orange) and MDAMB231 (red) on **(f)** PLL, **(g)** laminin, **(h)** fibronectin and **(i)** the uncoated electrode. The dashed line is the cell-free coated electrode. **(j)** Early slope of ΔR at 1 kHz (Ω h^-1^) for each line and substrate. All values are expressed relative to the coated electrode immediately before cell seeding. Full per-substrate four-channel comparisons are in *Figures S2–S4*, 17 h values and statistics in *Table S1*.

Because the coatings themselves modify the electrode interface, cell-free electrodes carrying each coating were monitored in DMEM for 120 h, including three medium changes visible as steps at 24, 48 and 70 h (**Figure 3a**). The protein-containing coatings, PLL+F and PLL+L in particular, show the largest capacitance increase, and PLL+F also yields the largest resistance decrease at 1 kHz. The bare electrode in medium shows the smallest capacitance change and the smallest resistance drift at 1 kHz, although at 41.5 kHz its resistance drifts as much as the coated electrodes over the full recording. Over the 17 h window the resistance drift is 7–25 Ω at 1 kHz (smallest on the uncoated electrode, largest on PLL+F) and 1–3 Ω at 41.5 kHz, whereas the capacitance drift is 5–43 nF at both frequencies, again largest for PLL+F and PLL+L (**Table S1**). The cell-free traces are therefore shown as a reference in **Figure 3f–i** and **Figures S2–S4**. The drift is positive in capacitance at both frequencies and negative in resistance, i.e. opposite in sign to the cell-induced changes in ΔC at 41.5 kHz and in ΔR at both frequencies, but of the same sign as the cell-induced increase in ΔC at 1 kHz. MCF10A cells show the strongest coupling of the three lines (**Figure 3b**). At 1 kHz, ΔR rises steeply over the first 10 h (**Figure 3e**). Fibronectin and PLL+F rise earliest and approach a plateau after 12 h, whereas PLL, PLL+L and the uncoated electrode are still rising at 17 h. The 41.5 kHz capacitance falls as the electrode becomes covered (**Figure 3b**). The decline is fastest and deepest on fibronectin and PLL+F, where ΔC reaches −71 and −73 nF, shallowest on laminin and the uncoated electrode (−56 and −59 nF), and intermediate on PLL (−62 nF) and PLL+L (−67 nF). The complementary channels (**Figure S1**) show the same picture: ΔC at 1 kHz rises sharply within the first 1–2 h and then more slowly, reaching 64–80 nF on the coated electrodes against 49 nF on the uncoated one. ΔR at 41.5 kHz rises sigmoidally, consistent with a spreading front that accelerates and then saturates as the layer closes. Fibronectin and PLL+F bring the cells to a coupled state fastest, but the coating also sets the endpoint: at 17 h, ΔC at 41.5 kHz and ΔR at both frequencies differ significantly across surfaces (p < 0.001; **Table S1**).

The spread across coatings in MCF10A is comparable to the gap between MCF10A and MDAMB468 on any given surface, and even to the gap between MCF10A and MDAMB231 on PLL and PLL+F. Only on the PLL-free surfaces is the MCF10A–MDAMB231 gap much larger than the coating effect (**Figure 3e, Table S1**). The intermediate MDAMB468 line shows rapid attachment: ΔC at 41.5 kHz drops by 20–30 nF within the first 2 h, and ΔR at 1 kHz reaches 50–80 Ω within 5 h with early slopes (13–18 Ω h^-1^) comparable to MCF10A (**Figure 3c, j**), after which both channels flatten. ΔR at 1 kHz plateaus at 67–87 Ω (**Figure 3e**). ΔC at 1 kHz grows slowly after the first 3 h and ends at 18 nF on the uncoated electrode, 26–27 nF on PLL and laminin, 35 nF on fibronectin and 40 nF on PLL+F and PLL+L (**Figure 3f–i, Figure S1c**). ΔR at 41.5 kHz rises only to 11–15 Ω over the full window (**Figure S1c**). The early plateau indicates that MDAMB468 cells attach and spread quickly but do not enhance coupling further. On fibronectin the 41.5 kHz capacitance falls least, albeit with large variability (±30 nF), and the 1 kHz resistance is lowest (67 Ω). Fibronectin therefore yields the least electrode coverage in this line, the opposite of its effect in MCF10A. Conversely, PLL+L yields the deepest capacitance drop, and PLL the highest 1 kHz resistance. However, none of the cell-dominated channels (ΔR at either frequency, ΔC at 41.5 kHz) differs significantly across coatings at 17 h (**Table S1**). The only significant channel is ΔC at 1 kHz.

The invasive line MDAMB231 gives the weakest electrical signals and the ones most shaped by substrate (**Figure 3d**). At 1 kHz, ΔR rises approximately linearly with no sign of saturation, and even the strongest condition (94 Ω on PLL) ends below the MCF10A range (103–137 Ω, **Figure 3e**). On PLL and PLL+F, ΔR reaches 90 Ω, on fibronectin and PLL+L 44–48 Ω, on laminin and the uncoated electrode only 15–23 Ω, comparable in magnitude to the cell-free coating drift over the same window (**Figure 3a**). Early slopes are correspondingly low, 1–6 Ω h^-1^ against 13–24 Ω h^-1^ for the other two lines (**Figure 3j**). The 41.5 kHz capacitance declines steadily, indicating a slowly covering layer. The decline is steepest on PLL+F (−71 nF), followed by PLL (−60 nF) and fibronectin (−48 nF), whereas PLL+L, the uncoated electrode and laminin reach only −31, −27 and −22 nF, less than half the drop recorded for MCF10A on the same surfaces. Likewise, ΔR at 41.5 kHz spans 11 Ω on the uncoated electrode, laminin and PLL+L to 30 Ω on PLL+F (**Figure S1d**). The kinetics represent a population of individually attached cells that spread and proliferate without forming a continuous electrically tight layer, which is the expected behavior of a mesenchymal-like line^27^. PLL alone and PLL under fibronectin drive the fastest and strongest initial coupling, ahead of fibronectin, whereas PLL under laminin does not: on PLL+L the 41.5 kHz capacitance behaves like laminin and the uncoated electrode, and the 1 kHz resistance like fibronectin, suggesting that the acceleration depends on the overlying protein.

### Spatial mapping of steady-state impedance

Cell adhesion was mapped across PEDOT:PSS microelectrode arrays sharing materials and experimental setup, providing complementary information (**Figure 4**). The MCF10A arrays show a lattice of high-resistance spots coinciding with the electrode features. The MDAMB468 arrays occupy the middle of the scale, with higher |Z| on PLL+L and laminin (1.03–1.08 × the uncoated value; **Figure 4b**, middle) and by far the highest spatial heterogeneity (CV, intended as the ratio of the standard deviation to the mean,, of 15–21% against 4–9% for the other two lines), consistent with growth in cohesive islands rather than a continuous layer. The MDAMB231 arrays sit at the bottom of the scale (**Figure 4a**). **Figure 4b (**left) shows the average values, with MCF10A between 40 and 52 kΩ, MDAMB468 between 35 and 42 kΩ and MDAMB231 between 31 and 41 kΩ. The comparison across cell lines and coatings indicates morphology and layer organization, whereas comparisons across coatings within the same line are informative on adhesion. In both MCF10A and MDAMB231 the uncoated electrode gives the highest resistance, 52 kΩ and 41 kΩ respectively, and fibronectin the second highest. In MCF10A, PLL and laminin give the lowest values, with PLL+F and PLL+L at ∼43 kΩ. In MDAMB231, PLL and PLL+F give the lowest values, with laminin, PLL+L and fibronectin at 34–36 kΩ. Note that this ordering differs from the 17 h kinetics, in which fibronectin and PLL+F gave the largest ΔR and PLL matched the uncoated electrode (**Figure 3e**). Because the 24 h maps are not referenced to the coated, cell-free electrode, part of the reduction on coated surfaces reflects the lower intrinsic impedance of the coatings themselves (**Figure 3a**). In MDAMB468, the laminin-based coatings give the highest resistance, while fibronectin, PLL and PLL+F are lowest at 35 kΩ. This matches the kinetics for this line, where PLL+L supported the deepest coverage and fibronectin the lowest. For MDAMB231, spatial maps and kinetics point in opposite directions. The PLL and PLL+F surfaces that drove the fastest and largest resistance rise during attachment are the surfaces with the lowest |Z| at 24 h (∼0.76 × the uncoated value), with no gain in spatial uniformity relative to the other coatings (**Figure 4b**, right). This indicate that the rapid coupling of MDAMB231 cells onto PLL does not translate into a high-resistance layer. The migration data shown later provide a functional context.

**Figure 4.**
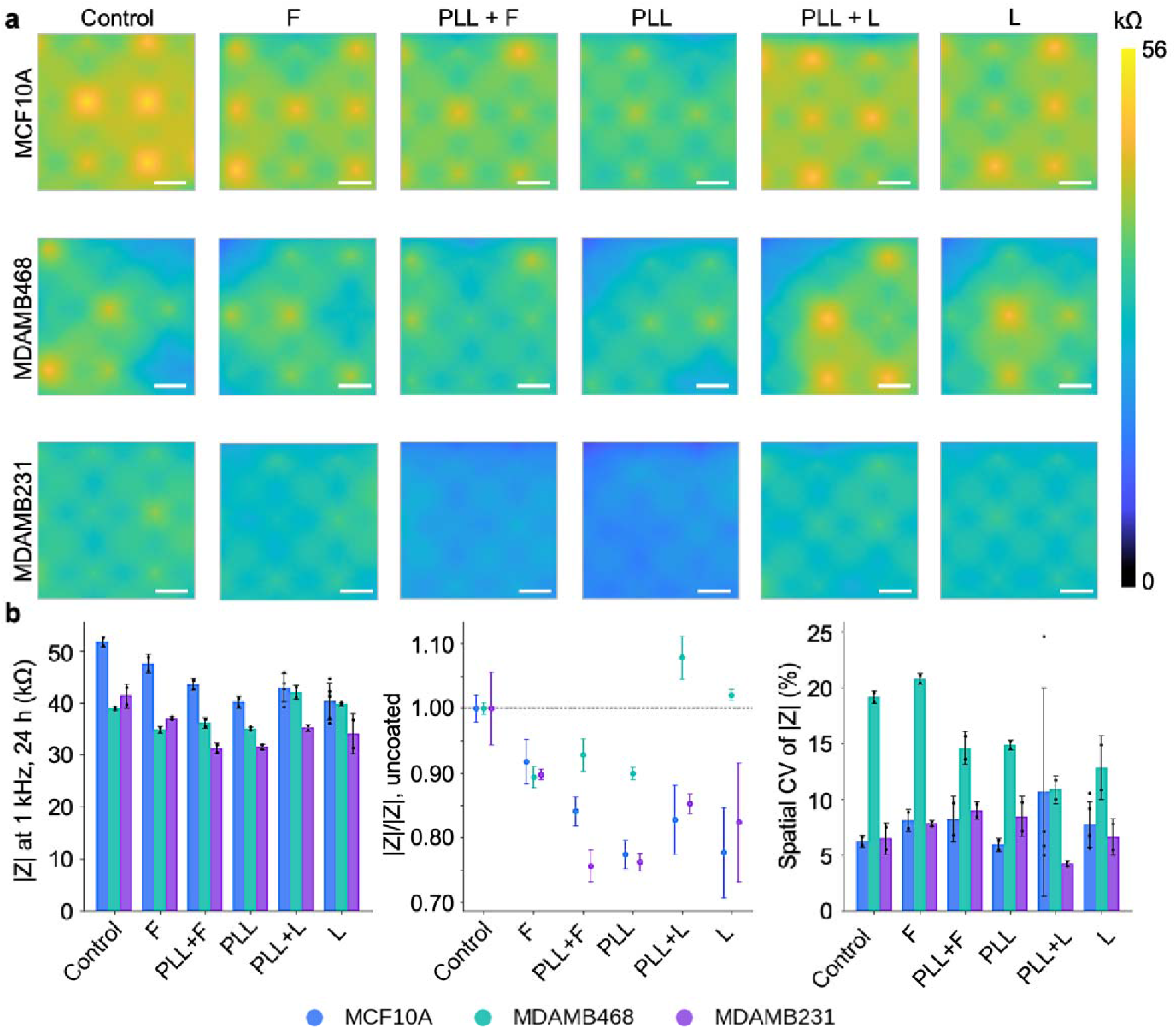
Spatial mapping of steady-state impedance at 24 h. **(a)** Maps of |Z| at 1 kHz across the microelectrode arrays 24 h after seeding MCF10A (top), MDAMB468 (middle) and MDAMB231 (bottom) cells on the six substrates. Regular lattices of elevated |Z| coincide with the electrode features and indicate uniform coverage, irregular patches indicate island-like growth. Scale bars: 200 µm. **(b)** Left: mean |Z| at 1 kHz per condition, grouped by cell line. Middle: the same values normalized to the uncoated electrode of each line. Right: spatial coefficient of variation of |Z| across each array, a measure of coverage heterogeneity. Bars/points are mean ± SD (n = 3 arrays per condition).

### Coating-dependent migration kinetics

The functional consequence of each cell–substrate interaction was examined for the two extremes of the spectrum, i.e. MCF10A and MDAMB231, in 50 h migration assays and origin-aligned single-cell trajectories (**Figure 5**). In MCF10A, closure proceeds steadily for the first 25–30 h and then levels off, with the final extent set by the substrate (**Figure 5a, Supplementary video S1**). The uncoated electrode supports the fastest and most complete closure in the non-tumorigenic line, reaching ∼90% at 50 h. Fibronectin produces a faster start than the remaining coatings, with 55% closure by 15 h, but plateaus at 80%. PLL, laminin and PLL+L settle at 70–75%. PLL+F is the slowest, reaching only ∼60% at 50 h. The curves share the same saturating form, which is consistent with epithelial collective migration^28^. Substrate chemistry therefore tunes the extent to which the MCF10A sheet migrates, but it does not change the migration mode. Fibronectin, which gave the fastest electrode coverage in *Figure 3b*, is among the fastest migrators. PLL+F, which coupled almost as strongly, is the slowest. Laminin and PLL+L, which coupled least strongly, result in migration at an intermediate rate. In this line, the strength of the initial cell–electrode coupling is unrelated to the speed of collective migration.

**Figure 5.**
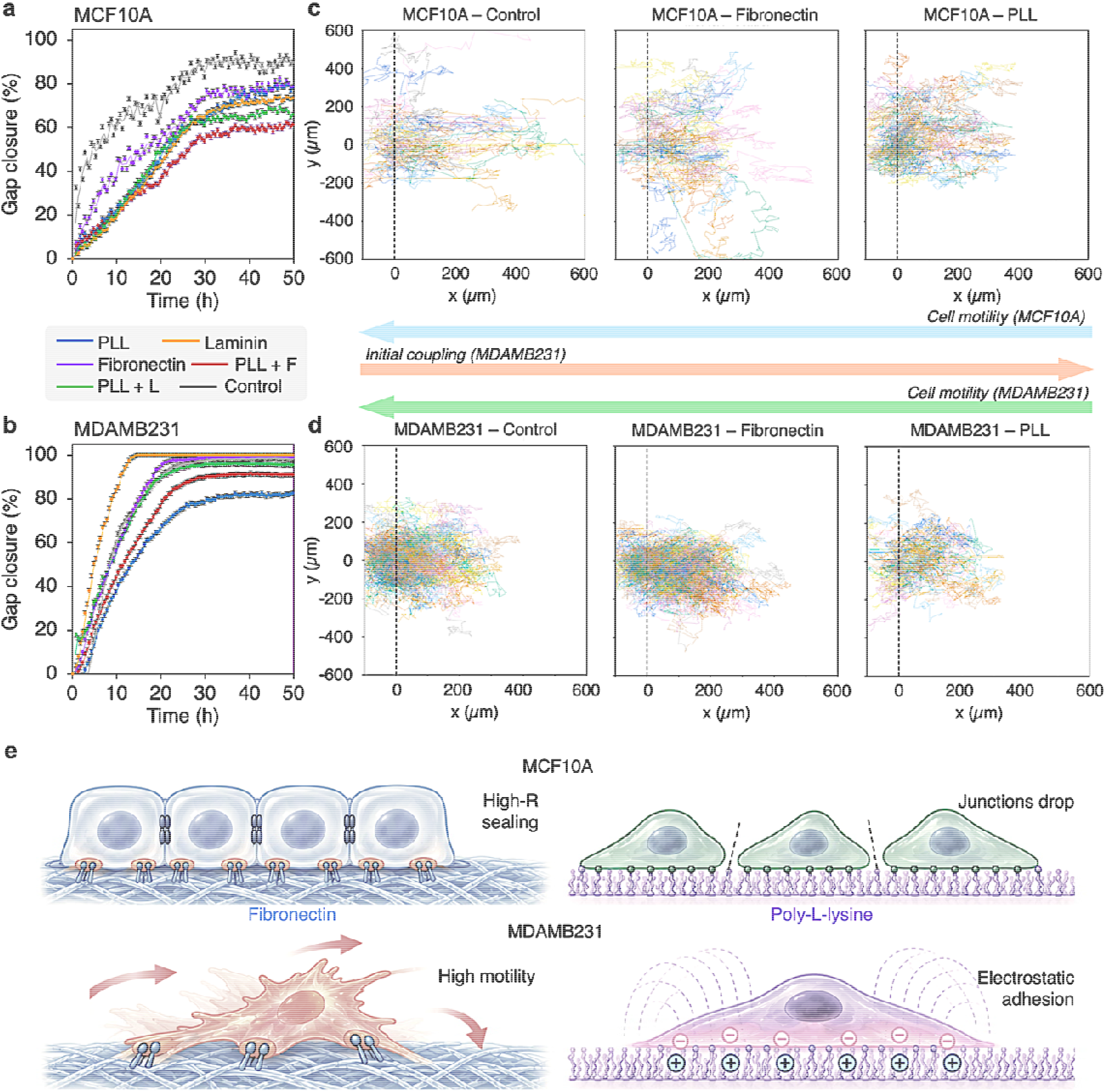
Coating-dependent migration, single-cell trajectories and proposed adhesion mechanisms. **(a, b)** Gap closure over 50 h for **(a)** MCF10A and **(b)** MDAMB231 cells on the six substrates. Data are mean ± SD (n = 3 wells per condition). **(c, d)** Origin-aligned single-cell trajectories of **(c)** MCF10A and **(d)** MDAMB231 cells at the wound edge on three representative substrates, ordered from the uncoated electrode through a physiological ligand to the synthetic polycation: control, fibronectin and PLL. Each line is one cell. x = 0 (dashed line) marks the initial position of each cell and positive x the direction of closure. n = 200 cells per panel, tracked for the entire migration assay, so track lengths should be compared within rather than across lines. Horizontal arrows between (c) and (d) indicate the direction of increasing cell motility (MCF10A and MDAMB231) and of increasing initial coupling at 17 h (MDAMB231) across the three substrates. In MCF10A initial coupling is not monotonic across these surfaces (fibronectin > PLL ≈ control) and is therefore not represented visually. **(e)** Proposed biophysical interpretation summarizing Figures 2–5. On fibronectin (left), MCF10A might form a junction-rich high-resistance epithelial sheet, whereas MDAMB231 remain individually motile possibly through integrin-mediated adhesion. On poly-L-lysine (right), MCF10A form a layer of lower steady-state resistance and more confined motility, consistent with reduced junctional organization, and MDAMB231 might be held by charge-mediated electrostatic adhesion that produces strong early coupling without a high-resistance interface.

In the MDAMB231 line, closure is faster (**Figure 5b, Supplementary video S2**). Laminin supports the most rapid closure, reaching 100% by 13 h. Fibronectin and the uncoated electrode reach 100% by 18–20 h. The three PLL-containing surfaces are the three slowest: PLL+L plateaus at about 96%, PLL+F at about 90% and PLL alone at about 83%. PLL is further distinguished by a lag of ∼3 h before any closure begins, during which all other MDAMB231 conditions are already advancing. Across the six surfaces the spread in closure kinetics is far wider than in MCF10A, and every PLL-free surface outperforms every PLL-containing one. The two surfaces on which MDAMB231 cells coupled fastest and most strongly, PLL and PLL+F, are the two on which they migrate slowest. The two surfaces on which they coupled most weakly, laminin and the uncoated electrode, are among the three on which they migrate fastest. To see how the collective closure kinetics arise from individual movement, cells at the wound edge were tracked and their paths plotted from a common origin at the edge (x = 0) for three representative surfaces, ordered from the bare electrode through a physiological ligand to the synthetic polycation: control, fibronectin and PLL. The two lines move in visibly different ways. MCF10A tracks (**Figure 5c**) spread widely in y, perpendicular to the direction of closure. On the uncoated electrode they advance furthest, on fibronectin they advance less while spreading across the full range in y. On PLL the cloud is more compact, with few tracks beyond about 380 µm in x. MDAMB231 tracks (**Figure 5d**) form dense clouds confined to ±250 µm in y and advancing 450 µm into the gap on both control and fibronectin. On PLL the cloud is shorter, with most tracks ending within about 330 µm.

## Discussion

This study focuses on how electrode surface chemistry changes the recorded electric signal of breast cells of different malignant potential, and how this relates with phenotype. This is relevant not only for the specific type of recordings adopted herein, but also as a reference for interface selection in cancer bioelectronics. The non-tumorigenic MCF10A line was the most strongly coupled and the least sensitive to coating, the invasive MDAMB231 line was the most weakly coupled and the most sensitive, and the intermediate MDAMB468 line coupled quickly and plateaued. The most informative outcome is that the relationship between coupling and motility is inverse in MDAMB231 and absent in MCF10A. In MDAMB231, the surfaces that drove the fastest and largest impedance rise after seeding, PLL and PLL+F (**Figure 3d**), gave the lowest |Z| at 24 h with no gain in spatial uniformity (**Figure 4**), the slowest gap closure (**Figure 5b**) and the most compact trajectories (**Figure 5d**), whereas the weakest-coupling surfaces, laminin and the uncoated electrode, supported the fastest migration. A strong early impedance signal in this phenotype might reflect cells captured and held at the surface rather than a functional interface, and the 3 h lag before PLL-seeded cells begin to move (**Figure 5b**) suggests they might first require loosening a non-physiological grip.

In MCF10A no such relationship exists: fibronectin and PLL+F produced near-identical rapid coupling (**Figure 3b**) yet sat at opposite ends of the migration range (**Figure 5a**). Laminin and PLL+L coupled least, whereas PLL coupled as strongly as the uncoated electrode, yet all three were indistinguishable in closure (**Figure 5a**) and PLL and laminin gave the same 24 h |Z| (**Figure 4b**). The trajectories explain the difference. MCF10A tracks wander perpendicular to the direction of closure (**Figure 5c**), suggesting that migration is collective and that the substrate can modulate how far the sheet advances but not how it advances. **Figure 5e** summarizes the interpretation. On fibronectin, MCF10A form a high-resistance sheet while MDAMB231 remain individually motile. On PLL, MCF10A form a less organized layer and MDAMB231 are held by a charge-mediated adhesion that neither matures into a well-coupled layer nor enhances migration. PLL deserves separate comment because it is the surface most often used in cell–electrode interfaces^29-31^. It presents a dense positive charge to a negatively charged membrane and promotes adhesion without engaging integrins^17^. In MCF10A it lowered 24 h resistance relative to the uncoated electrode (**Figure 4b**) and gave the most compact trajectories (**Figure 5c**), so even in a junction-forming epithelium the cationic surface interferes with the organization of the layer. Layering fibronectin on PLL did not rescue either cell line: PLL+F was the slowest migrator in MCF10A and the second slowest in MDAMB231. Therefore, PLL reduces motility. The 17 h kinetics and the 24 h maps were acquired on different arrays and with different referencing, and the coating drift was not subtracted from the cell traces. The cell-free references are provided so that readers can judge which channels are cell-dominated.

## Conclusion

The electrode coating is an experimental variable that interacts with phenotype and carries intrinsic electrical signature. Uncoated electrodes and fibronectin-coated electrodes preserved the largest differences between the non-tumorigenic and invasive lines in both impedance and motility, whereas PLL compressed those differences by limiting MDAMB231 motility and coupling. The same property that makes PLL unsuitable for unbiased phenotyping could, conversely, be exploited: a surface that selectively limits invasive cell migration is a potential tool for studying adhesion-dependent drug responses. Two practical consequences follow. First, the same cells on different coatings can differ in impedance as much as different cells on the same coating. Second, the coating itself can be made part of the readout: patterning several chemistries on a single array would yield a multi-surface impedance fingerprint in which the spread of responses reports on how dependent a population is on cell– substrate adhesion. Future work should focus on analyzing the interface and adhesion with molecular tools, and on expanding the analysis to other cell types to identify broader trends, their mechanisms and implications for *in vitro* biophysical and translational studies targeting cancer bioelectricity.

## Materials and Methods

### Cell culture

MDA-MB-231 and MDA-MB-468 cells were grown in high-glucose DMEM with 10% FBS, 1% L-glutamine and 1% penicillin/streptomycin. MCF10A cells were grown in DMEM/F12 with 5% horse serum, 500 ng/mL hydrocortisone, 10 µg/mL insulin and 20 ng/mL EGF. All lines were kept at 37 °C, 5% CO_2_, medium was replaced three times per week, and cells were passaged at 80% confluence.

### Electrode platforms

Two Axion Biosystems (USA) platforms were used. (i) Time-resolved coupling (**Figure 3**) and migration (**Figure 5**) were performed on CytoView[Z 96-well plates, in which each well carries a single gold recording electrode embedded in a transparent PET bottom (100 µm finger width, 100 µm gap, ∼20 mm^2^ sensing area), read on a Maestro Z system with AxIS Navigator (v3.12.7.2). (ii) Spatial resistance mapping (**Figure 4**) was performed on CytoView MEA 24-well plates, in which each well carries 16 PEDOT-coated gold microelectrodes (50 µm diameter) in a 4×4 grid with 350 µm pitch (1.1×1.1 mm recording area), read on a Maestro Edge system with the MEA Viability module. Both systems were kept at 37 °C and 5% CO_2_ during recording.

### Coatings

Fibronectin and laminin stocks (1 mg/mL in dH_2_O) and poly-L-lysine (PLL, 0.01%) were aliquoted and stored at 4 °C. Coating densities are given as µg/cm^2^. For CytoView[Z wells (0.32 cm^2^, 50 µL) C(µg/mL) = 6.4×C(µg/cm^2^). 3 µg/cm^2^ fibronectin and 1 µg/cm^2^ laminin were therefore prepared as 19 µg/mL and 6.4 µg/mL working solutions in PBS and applied as 50 µL per CytoView[Z well or as an 8 µL drop over the electrode area of CytoView MEA wells. PLL (same volumes) was incubated >45 min at room temperature, washed three times with dH_2_O and air-dried. For PLL+F and PLL+L, protein solution was then added onto the PLL layer and incubated overnight at 4 °C. Immediately before seeding, excess liquid was aspirated without washing and wells were not allowed to dry. Uncoated wells served as controls. Six conditions were tested: uncoated, PLL, fibronectin, laminin, PLL+F and PLL+L.

### Live/dead assay

Cells were seeded on coated CytoView[Z wells at 25000 cells/well and stained 24 h later. After a gentle PBS wash, cells were incubated 30 min in PBS containing 1 µM calcein[AM (live, green) and 5 µM ethidium homodimer[1 (dead, red) and imaged in the green and red fluorescence channels of an Omni live-cell imager (37 °C, 5% CO_2_). Live and dead fractions were counted in n = 3 fields per condition.

### Time-resolved impedance

Coated CytoView[Z 96-well plates received 100 µL of medium per well and were docked on the Maestro Z to record the coated-electrode reference. Then 100 µL of cell suspension was added (25000 cells/well, 200 µL final, three wells per condition) and resistance and capacitance were recorded at 1 kHz and 41.5 kHz every 15 min for 17 h. All traces are expressed as changes (ΔR, Ω; ΔC, nF) relative to the coated cell-free electrode immediately before seeding. Cell-free coated electrodes were recorded in the same way in DMEM for 120 h, with medium renewals at 24, 48 and 70 h, to characterize the intrinsic coating signal. Endpoint values at 17 h were compared across substrates by one-way ANOVA and Kruskal–Wallis tests (**Table S1**). Early slopes (**Figure 3j**) were obtained by linear regression of ΔR at 1 kHz over after seeding.

### Spatial resistance mapping

Coated CytoView MEA 24-well plates were seeded with 15000 cells/well as an 8 µL droplet over the electrode area, and 0.5 mL of medium was added after 2 h. At 24 h, per-electrode resistance at 1 kHz (kΩ) was acquired in the viability (impedance) mode of AxIS Navigator and exported with the AxIS Statistics Compiler. Electrodes with resistance ≥18 kΩ (software default) were classed as cell-covered. Maps were generated by interpolation of the 16 electrode values over the array. For each array, the mean |Z| and the spatial coefficient of variation CV (%) = (σ/µ) × 100 across electrodes were computed with n = 3 arrays per condition.

### Migration assay

Cells were seeded at 25000 cells/well in coated CytoView[Z 96-well plates and grown to confluence. A linear scratch was made with a p10 tip, wells were washed with PBS and refilled with 200 µL of medium. Whole-well brightfield images (2.63 µm/pixel) were acquired every 30 min for 50 h on the Omni system (37 °C, 5% CO_2_). Wound area was quantified with a custom Python pipeline (OpenCV, NumPy) applied identically to all conditions. Pixel size was calibrated from the embedded scale bar, and analysis was restricted to the electrode-free central region. Stage drift was corrected by phase-correlation alignment of each frame to the electrodes, and cell-free regions were identified by low local edge density. An Otsu threshold set between the fresh scratch and the intact monolayer at 0 h was corrected for a small temporal drift measured in cell-free regions. Patches <2000 µm^2^ were discarded. The wound was defined as the image columns cell-free over ≥50% of the field height at 0 h, extended by ∼50 µm on each side. Closure = 100 × (1−A(t)/A_0_), with A(t) the cell-free wound area at time t and A_0_ that at the first usable frame. Mean ± SD of three wells per condition.

### Single-cell tracking

Cell migration video analysis. Time-lapse videos of migration assays were analyzed with Python 3.10.12 (including OpenCV, NumPy, SciPy, and Matplotlib), to extract single-cell trajectories. Each frame was converted to greyscale, smoothed with a 5×5 Gaussian kernel, and segmented by inverted adaptive Gaussian thresholding (block size 21 px, offset 4). External contours were extracted and retained as cells if their area fell between 150 and 350 px^2^, and the centroid of each retained contour was computed from its image moments. Cells were linked between consecutive frames by nearest-neighbor matching of centroids. Unmatched detections were initialized as new tracks, and tracks with no matching detection for more than 3 consecutive frames were discarded. Only tracks longer than 8 frames were kept for analysis, which excludes transient detections of debris and segmentation noise. For each retained track the coordinates were exported together with the displacement (Δx, Δy) from the cell first detected position. Origin-aligned plots were generated by translating each track so that its first position coincides with (0, 0), with the positive x direction oriented towards the gap.

