## Supplementary file for "Substrate chemistry in cancer bioelectronics: phenotype-dependent cell-electrode coupling and motility"

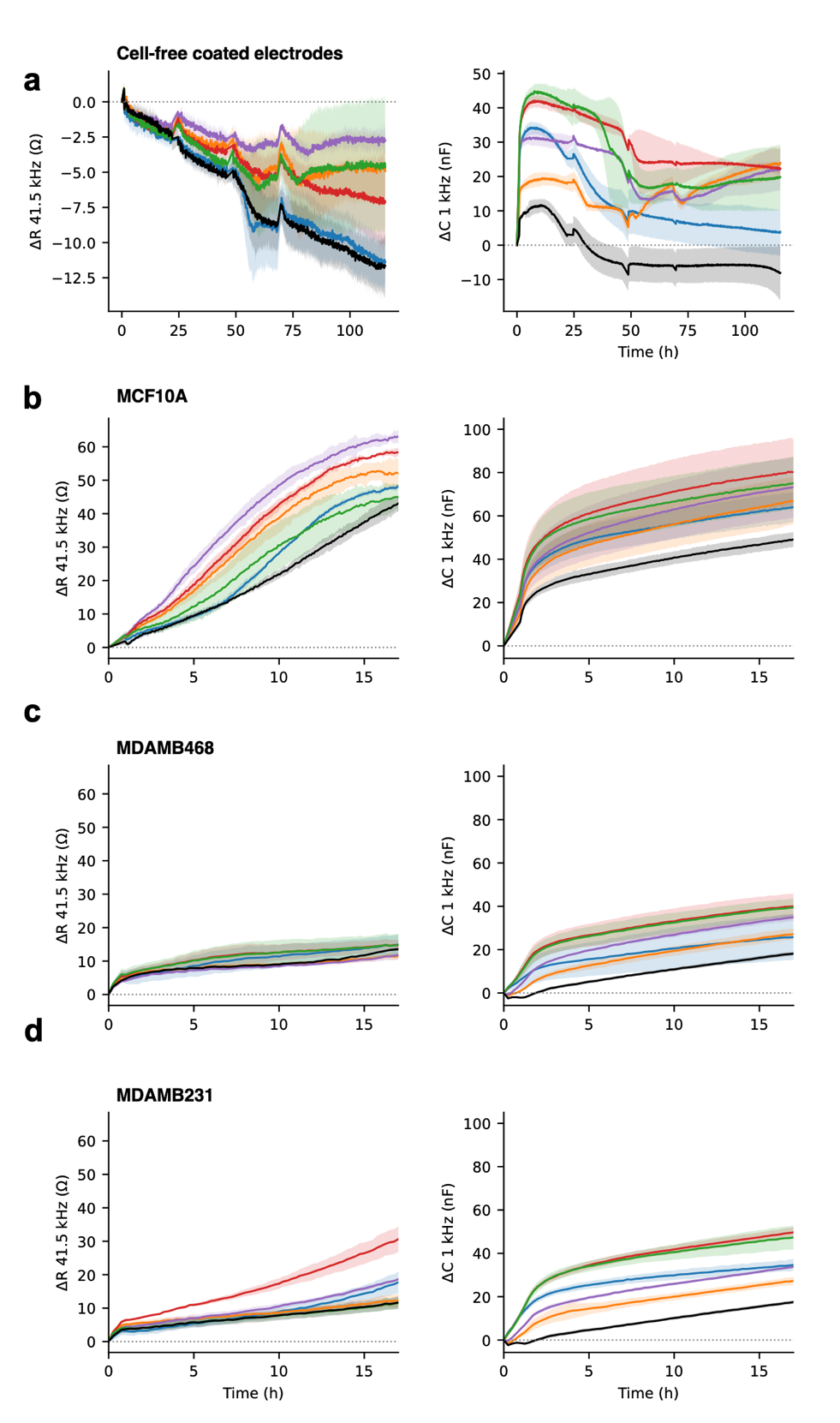

**Supplementary Figure S1.** Complementary impedance channels to Figure 3. ΔR at 41.5 kHz (left) and ΔC at 1 kHz (right) for **(a)** cell-free coated electrodes in DMEM over 120 h, and **(b)** MCF10A, **(c)** MDAMB468 and **(d)** MDAMB231 cells over the first 17 h after seeding. Colors as in Figure 3a–d: PLL (blue), laminin (orange), fibronectin (purple), PLL + F (red), PLL + L (green), uncoated (black). Mean ± SD.

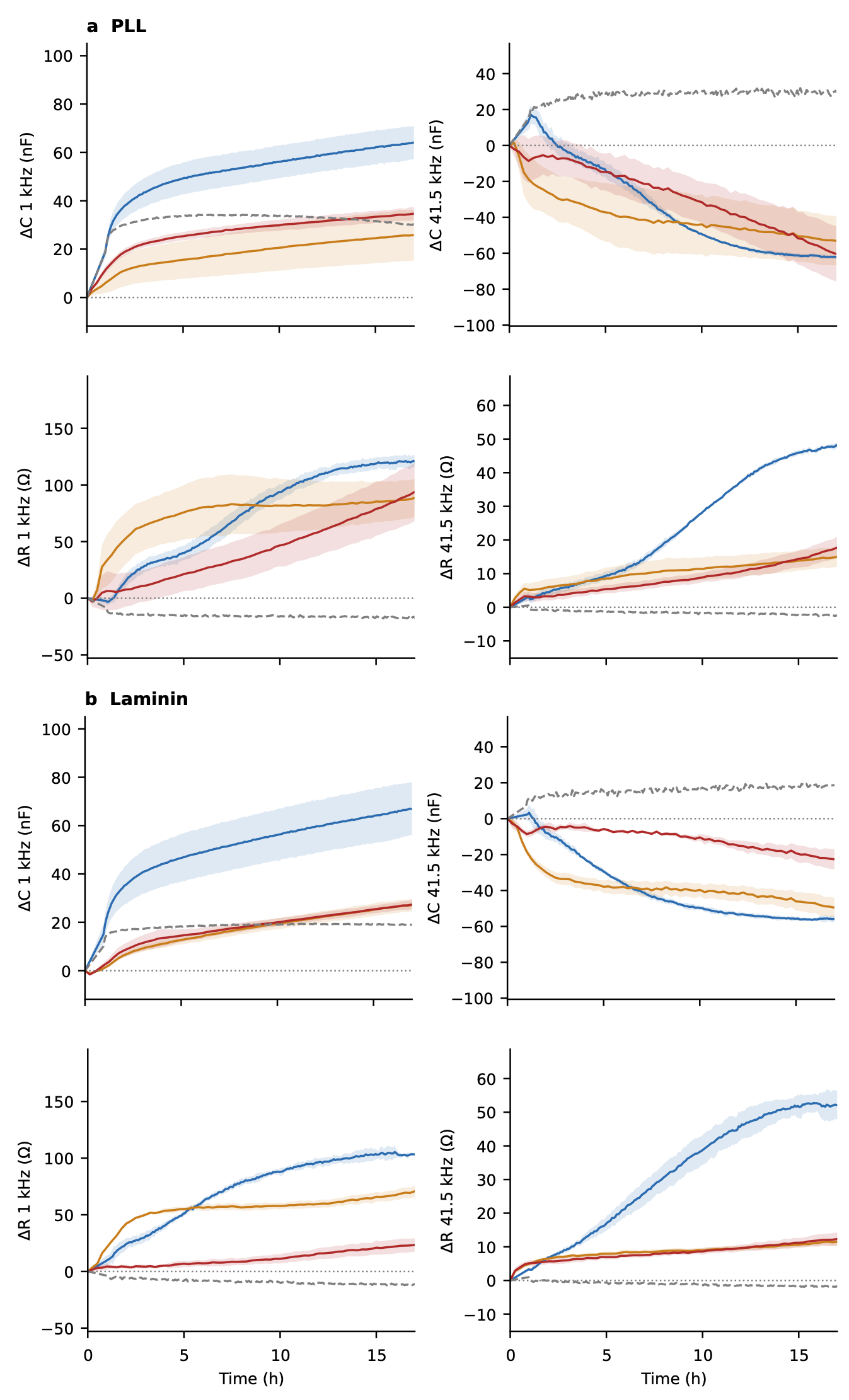

**Supplementary Figure S2.** Line-by-line comparison on **(a)** PLL and **(b)** laminin. For each substrate, ΔC at 1 kHz, ΔC at 41.5 kHz, ΔR at 1 kHz and ΔR at 41.5 kHz over 17 h for MCF10A (blue), MDAMB468 (orange) and MDAMB231 (red) are reported. Grey dashed line indicate cell-free coated electrode. Mean ± SD.

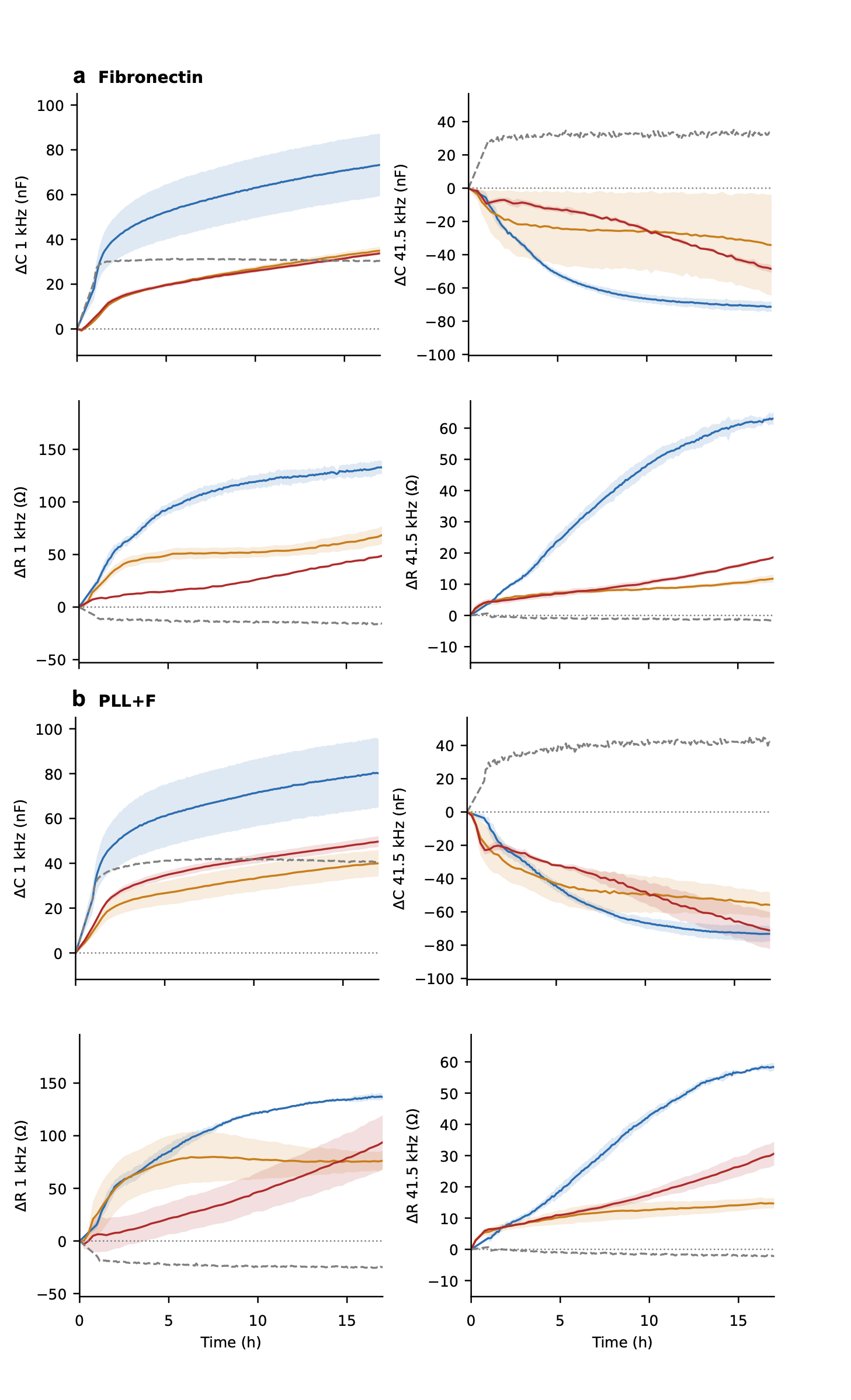

**Supplementary Figure S3.** Line-by-line comparison on **(a)** fibronectin and **(b)** PLL+fibronectin. For each substrate, ΔC at 1 kHz, ΔC at 41.5 kHz, ΔR at 1 kHz and ΔR at 41.5 kHz over 17 h for MCF10A (blue), MDAMB468 (orange) and MDAMB231 (red) are reported. Grey dashed line indicate cell-free coated electrode. Mean ± SD.

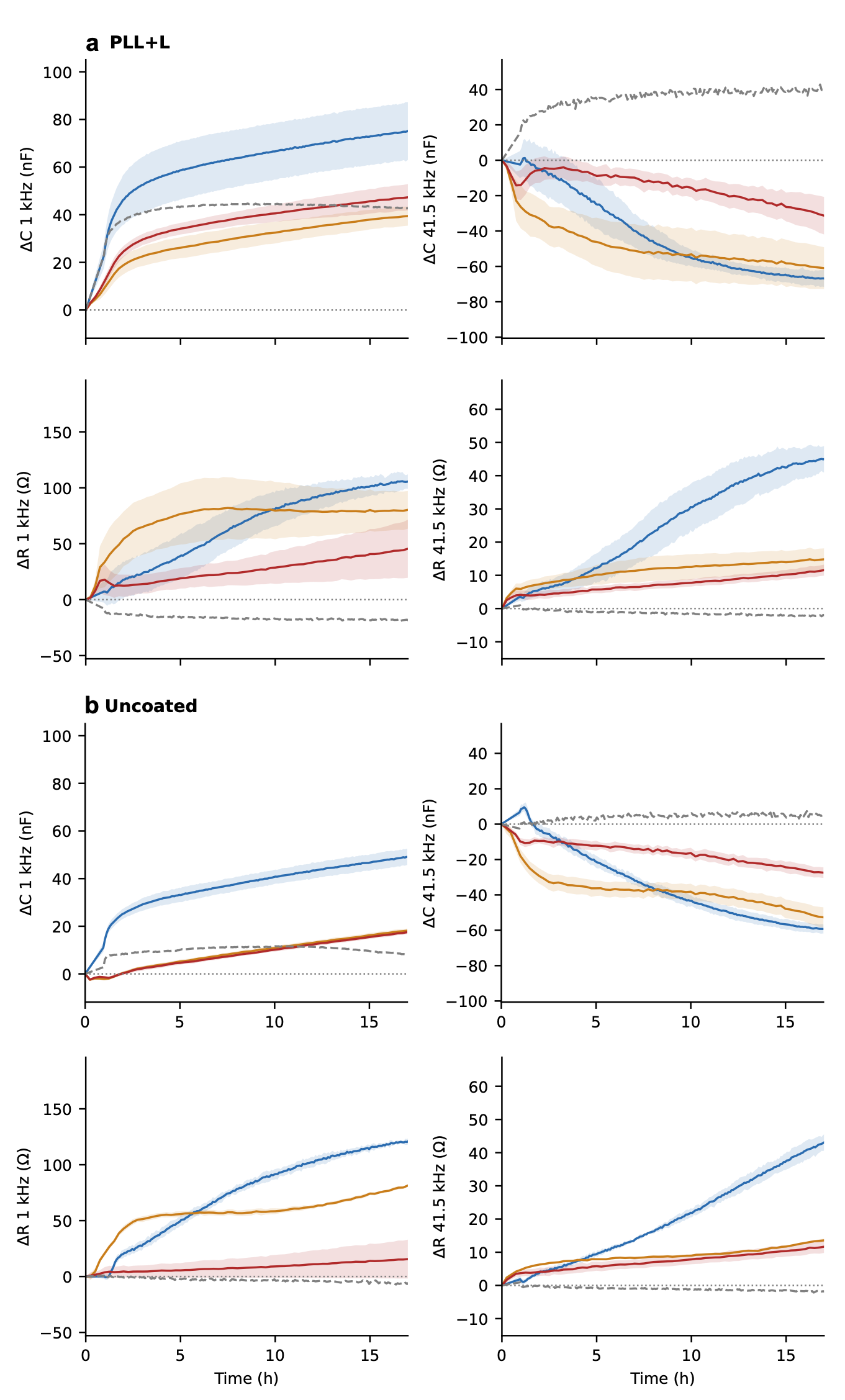

**Supplementary Figure S4.** Line-by-line comparison on **(a)** PLL+laminin and **(b)** uncoated electrode. For each substrate, ΔC at 1 kHz, ΔC at 41.5 kHz, ΔR at 1 kHz and ΔR at 41.5 kHz over 17 h for MCF10A (blue), MDAMB468 (orange) and MDAMB231 (red) are reported. Grey dashed line indicate cell-free coated electrode. Mean ± SD.

**Supplementary Table S1.** Changes at 17 h relative to the coated electrode before seeding (mean ± SD) for each line, channel and substrate, with ANOVA and Kruskal–Wallis p-values across substrates. Cell-free rows give the coating drift over the same 17 h. Capacitance is expressed in nF, absolute impedance in Ω.

| *Cell line* | *Channel* | *PLL* | *Laminin* | *Fibronectin* | *PLL+F* | *PLL+L* | *Uncoated* | *ANOVA p* | *Kruskal–Wallis p* |
| --- | --- | --- | --- | --- | --- | --- | --- | --- | --- |
| MCF10A | ΔC 1 kHz | 64.1 ± 6.8 | 66.9 ± 10.9 | 73.2 ± 14.0 | 80.2 ± 15.3 | 75.0 ± 12.2 | 49.1 ± 3.4 | n.s. | n.s. |
| MCF10A | ΔC 41.5 kHz | -62.1 ± 1.0 | -55.9 ± 1.9 | -71.4 ± 3.1 | -73.2 ± 4.5 | -66.8 ± 4.7 | -59.2 ± 2.7 | p < 0.001 | p = 0.016 |
| MCF10A | ΔR 1 kHz | 121.3 ± 5.4 | 103.4 ± 0.9 | 133.0 ± 6.2 | 136.8 ± 3.2 | 105.4 ± 6.2 | 120.3 ± 2.6 | p < 0.001 | p = 0.011 |
| MCF10A | ΔR 41.5 kHz | 48.1 ± 0.8 | 52.2 ± 4.3 | 63.1 ± 2.0 | 58.4 ± 1.3 | 44.9 ± 3.9 | 42.9 ± 2.4 | p < 0.001 | p = 0.008 |
| MDAMB468 | ΔC 1 kHz | 25.6 ± 10.5 | 26.9 ± 2.6 | 34.8 ± 1.7 | 39.8 ± 5.8 | 39.3 ± 4.1 | 17.9 ± 0.9 | p = 0.002 | p = 0.012 |
| MDAMB468 | ΔC 41.5 kHz | -52.8 ± 13.8 | -49.0 ± 5.5 | -33.9 ± 30.0 | -55.5 ± 7.8 | -60.7 ± 12.1 | -52.4 ± 5.6 | n.s. | n.s. |
| MDAMB468 | ΔR 1 kHz | 87.4 ± 16.8 | 69.3 ± 4.9 | 66.7 ± 8.1 | 75.5 ± 9.3 | 79.4 ± 17.2 | 79.6 ± 1.2 | n.s. | n.s. |
| MDAMB468 | ΔR 41.5 kHz | 14.7 ± 2.9 | 11.4 ± 1.1 | 11.6 ± 1.2 | 10.7 ± 1.6 | 14.7 ± 3.3 | 13.4 ± 0.5 | n.s. | n.s. |
| MDAMB231 | ΔC 1 kHz | 34.4 ± 2.8 | 27.0 ± 1.8 | 33.5 ± 0.5 | 49.4 ± 2.4 | 47.1 ± 5.4 | 17.2 ± 0.8 | p < 0.001 | p = 0.008 |
| MDAMB231 | ΔC 41.5 kHz | -59.5 ± 15.4 | -22.4 ± 5.6 | -47.8 ± 2.2 | -70.6 ± 11.0 | -30.6 ± 10.5 | -27.1 ± 3.0 | p < 0.001 | p = 0.015 |
| MDAMB231 | ΔR 1 kHz | 94.4 ± 30.1 | 22.9 ± 6.0 | 47.6 ± 1.3 | 91.3 ± 25.6 | 44.2 ± 25.2 | 15.2 ± 16.9 | p < 0.001 | p = 0.008 |
| MDAMB231 | ΔR 41.5 kHz | 17.1 ± 3.0 | 12.1 ± 1.9 | 18.1 ± 0.3 | 29.9 ± 3.7 | 11.3 ± 1.6 | 11.4 ± 1.9 | p < 0.001 | p = 0.016 |
| cell-free | ΔC 1 kHz | 30.3 ± 1.1 | 19.0 ± 1.7 | 30.5 ± 2.5 | 40.5 ± 2.0 | 42.7 ± 3.1 | 8.1 ± 2.3 | — | — |
| cell-free | ΔC 41.5 kHz | 30.4 ± 4.0 | 18.7 ± 0.1 | 33.3 ± 1.2 | 43.4 ± 1.7 | 39.9 ± 13.4 | 5.0 ± 3.5 | — | — |
| cell-free | ΔR 1 kHz | -16.4 ± 0.1 | -11.3 ± 1.3 | -15.7 ± 0.8 | -24.9 ± 3.4 | -17.9 ± 2.3 | -6.7 ± 1.4 | — | — |
| cell-free | ΔR 41.5 kHz | -2.5 ± 0.4 | -1.8 ± 0.3 | -1.4 ± 0.1 | -2.2 ± 0.1 | -1.9 ± 0.3 | -1.8 ± 0.2 | — | — |

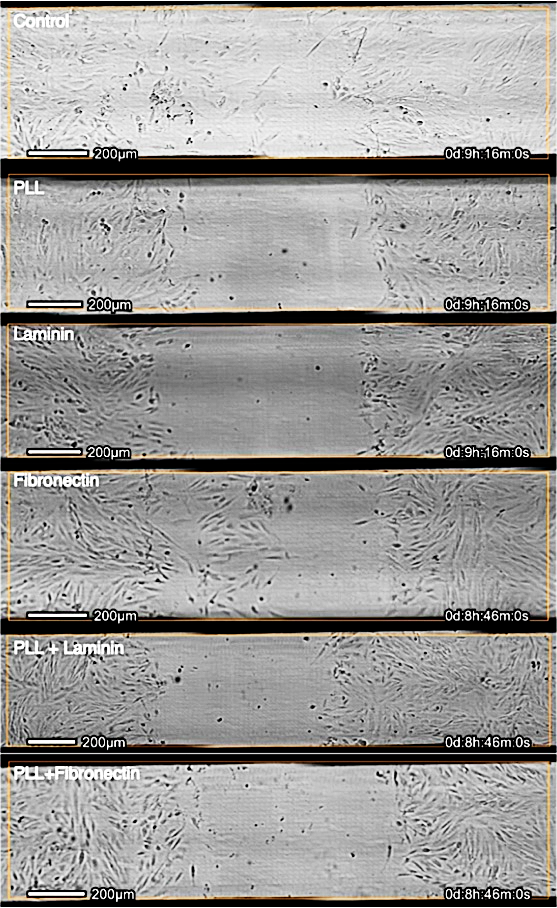

**Supplementary Video S1.** Migration of MCF10A in the six conditions.

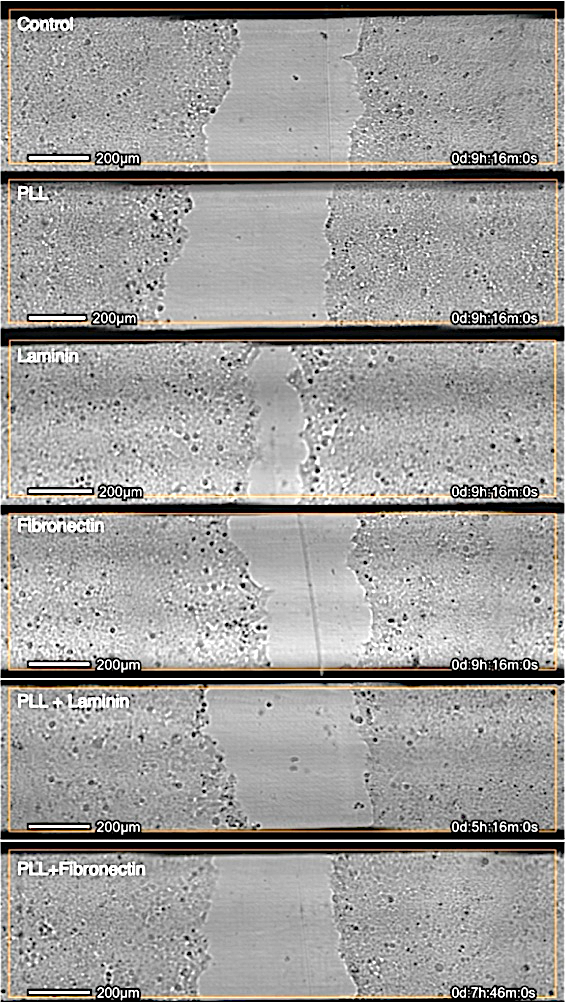

**Supplementary Video S2.** Migration of MDAMB231 in the six conditions.
